# Modeling the functional relationship network at the splice isoform level through heterogeneous data integration

**DOI:** 10.1101/001719

**Authors:** Hong-Dong Li, Rajasree Menon, Ridvan Eksi, Aysam Guerler, Yang Zhang, Gilbert S. Omenn, Yuanfang Guan

**Affiliations:** Department of Computational Medicine and Bioinformatics, University of Michigan, Ann Arbor, Michigan, United States; Department of Internal Medicine, University of Michigan, Ann Arbor, Michigan, United States; Department of Electrical Engineering and Computer Science, Ann Arbor, Michigan, United States

## Abstract

Functional relationship networks, which reveal the collaborative roles between genes, have significantly accelerated our understanding of gene functions and phenotypic relevance. However, establishing such networks for alternatively spliced isoforms remains a difficult, unaddressed problem due to the lack of systematic functional annotations at the isoform level, which renders most supervised learning methods difficult to be applied to isoforms. Here we describe a novel multiple instance learning-based probabilistic approach that integrates large-scale, heterogeneous genomic datasets, including RNA-seq, exon array, protein docking and pseudo-amino acid composition, for modeling a global functional relationship network at the isoform level in the mouse. Using this approach, we formulate a gene pair as a set of isoform pairs of potentially different properties. Through simulation and cross-validation studies, we showed the superior accuracy of our algorithm in revealing the isoform-level functional relationships. The local networks reveal functional diversity of the isoforms of the same gene, as demonstrated by both large-scale analyses and experimental and literature evidence for the disparate functions revealed for the isoforms of *Ptbp1* and *Anxa6* by our network. Our work can assist the understanding of the diversity of functions achieved by alternative splicing of a limited set of genes in mammalian genomes, and may shift the current gene-centered network prediction paradigm to the isoform level.

**Author summary:** Proteins carry out their functions through interacting with each other. Such interactions can be achieved through direct physical interactions, genetic interactions, or co-regulation. To summarize these interactions, researches have established functional relationship networks, in which each gene is represented as a node and the connections between the nodes represent how likely two genes work in the same biological process. Currently, these networks are established at the gene level only, while each gene, in mammalian systems, can be alternatively spliced into multiple isoforms that may have drastically different interaction partners. This information can be mined through integrating data that provide isoform-level information, such as RNA-seq and protein docking scores predicted from amino acid sequences. In this study, we developed a novel algorithm to integrate such data for predicting isoform-level functional relationship networks, which allows us to investigate the collaborative roles between genes at a high resolution.

## Introduction

Genes fulfill their functions by interacting with each other through complex biological networks. A key approach to systematically model such interactions is to establish functional relationship networks, which represent the probability of two proteins working in the same biological process. Such networks are generated through integrating heterogeneous, large-scale genomic datasets [1–25], such as gene expression profiles, transcription factor binding sites, protein-protein physical interaction and genetic interaction data. Diverse algorithms have been developed for modeling such networks, including Bayesian networks [2,5,7,8,26], support vector machines [27], the log-likelihood scoring scheme based on a Bayesian statistic approach [17,19–22,28]. Despite their differences, these data integration methods have the following common steps. First, a set of ground-truth functionally related gene pairs are defined by co-annotation to a specific biological function or a pathway [29–31]; these pairs constitute the gold-standard set. Then, diverse genomic datasets are collected. Finally, a model is constructed to leverage the informativeness of the genomic datasets using the gold-standard pairs and predict how likely two genes are working in the same biological process. These networks have shown great potential in providing new insights into gene functions and advancing our understanding of disease mechanisms [1,18,32,33].

However, the above pipeline cannot be readily extended to establishing networks at the isoform level due to two major obstacles. First, most of the traditional functional genomic data, such as most microarray expression and physical interaction, are routinely recorded or analyzed at the gene level, and thus do not directly provide isoform-level features. Protein domain datasets, on the other hand, are available at the isoform level, but cannot be used in many model organisms including mouse. This is because many functional annotations are directly mapped from protein domains, potentially leading to over-fitting in training our models. Fortunately, recent development of computational approaches and experimental technologies have provided multiple types of genomic data sources at the isoform level, including RNA-seq [34–37] and computationally predicted protein-protein docking scores [38]. The availability of these data has overcome the first obstacle facing network modeling at the isoform level.

The second challenge facing isoform-level network modeling remains: We do not have a set of ground-truth functionally related isoform pairs to serve as the gold standard to evaluate and integrate large-scale genomic data. Both biological functions [29] and pathways [30,31,39–41] are conventionally documented at the gene level but not at the isoform level, preventing any classical method used for building gene-level networks from being directly applied to splice isoforms.

In this paper, we proposed a novel conceptual framework to solve this problem, which is inspired by our recent work [42] where we developed an algorithm for predicting protein functions at the alternatively spliced isoform level. Our critical idea is treating a gene pair, for which we have a gold standard label, as a bag of multiple isoform pairs of potentially different probabilities to be functionally related. Our key problem here is how to identify the isoform pairs that are truly related, under the assumption that if a gene pair is functionally related, at least one of its isoform pairs must be related to allow co-functionality at the gene level; otherwise, none of its isoform pairs can be functionally related. The formulation of this problem falls into the category of multiple instance learning (MIL) [43–46]. In this paper, we tested several established MIL algorithms and developed a new Bayesian network-based MIL method in the context of network integration. Then, we established a genome-wide isoform-level network for the mouse through heterogeneous data integration. We showed that our isoform-level network can differentiate the connected and the unrelated isoform pairs of a single gene pair, providing a high-resolution view of the original gene-level network. Furthermore, we demonstrated that our isoform network reveals the functional diversity of the isoforms of the same gene. Our work is the first study that investigates functional relationships at the isoform level. We expect that this approach will advance the understanding of protein diversity achieved by alternative splicing and can be readily extended to other domains of computational predictions for isoforms.

## Results

### Formulating isoform-level network prediction into a multiple instance learning problem

As discussed in the Introduction, the key challenge facing isoform-level network modeling is the lack of ground-truth functionally related pairs. To solve this problem, we assume that: i) of a functionally related gene pair (a positive bag), at least one of its isoform pairs (the instances) must be functionally related (**Figure S1**); ii) for an unrelated gene pair (a negative bag), none of its isoform pairs can be functionally related. Then, our aim is to identify the truly functionally related isoform pairs (“witnesses”) of the positive bags. Under these two assumptions, our problem becomes a classical multiple instance learning (MIL) problem.

MIL is a widely-used and well-developed framework applicable to many domains where hidden labels for ‘instances’ need to be inferred while the bag labels are known. It has been used for predicting hand-written digits [47], small molecule shapes [45], drug interactions and image classifications [44,48]. MIL can be used to assign instance-level labels as well as to improve bag level predictions [44]. For example, in predicting whether a drug can bind to certain molecule, the drug is considered as a ‘bag’ of potential conformations (instances), and MIL can be used to predict whether the drug can bind to the molecule and find the conformations that bind (the witnesses). Recently, we also introduced this concept into the functional genomics field for predicting isoform functions [49]. MIL can be integrated into many base learners, such as support vector machines (SVM) [43], Bayesian classifiers [50] and decision trees [51]. Because our genomic datasets are characterized by large scale, heterogeneity, as well as missing data in some of the datasets, we focused on implementing established and novel MIL algorithms together with the Bayesian network (**Figure 1**). Bayesian networks have been most widely used in predicting gene-level functional relationship networks, including our previous gene-level functional relationship networks [1,2].

**Figure 1.**
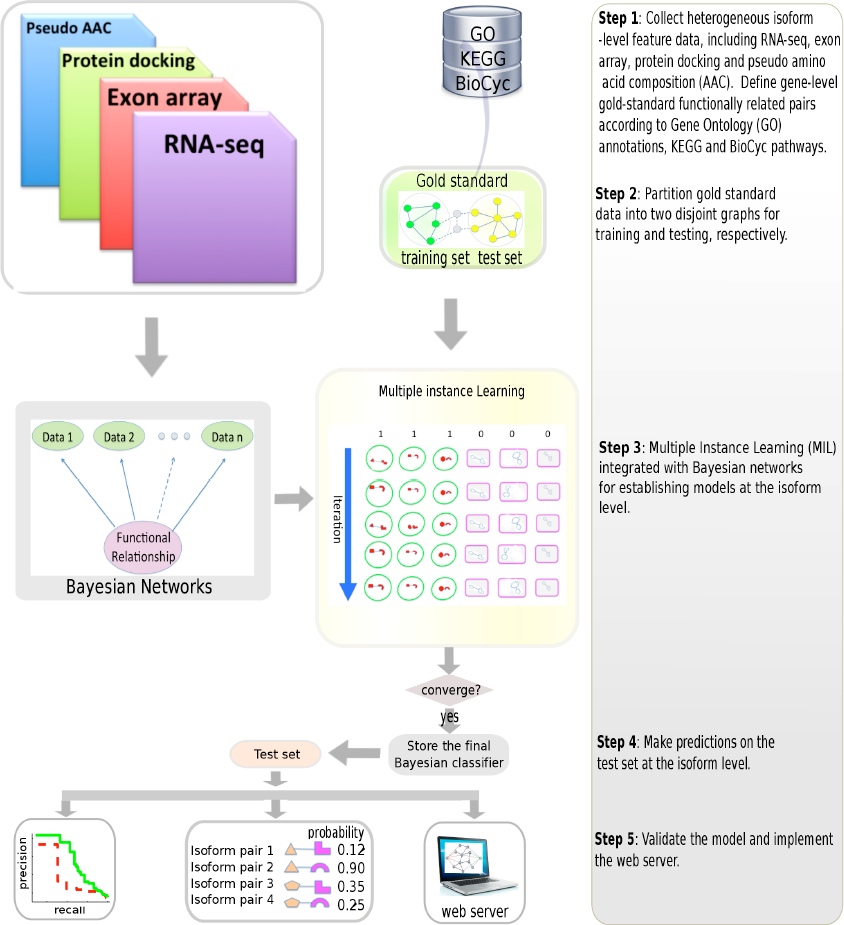
Modeling the isoform-level functional relationship network using Bayesian network-based multiple instance learning. We first collected genomic features from different data sources including RNA-seq, exon array, protein docking and pseudo amino acid compositions. For each dataset, pair-wise values for isoform pairs were calculated. We generated the gene-level gold standard, which contains positive gene pairs (co-annotated to the same biological function or pathway) and negative gene pairs (not co-annotated to any function/pathway) using the Gene Ontology (GO), KEGG and BioCyc databases. For model development and validation, we partitioned our gold standard into two disjoint graphs, serving as the training set and the test set, respectively. Our MIL algorithm, which uses a Bayesian network classifier as its base learner, was run on the training data to build a classification model. In each iteration, only the “witnesses” (red-colored pairs) in positive bags and the highest scored instance in negative bags are used for training the model to achieve maximal discriminativeness. Therefore, the classification model was established at the isoform level, instead of at the gene level. After convergence, the final classifier was used to predict the probability of functional interactions for the independent test set. We finally validated the accuracy of our model through simulation, cross-validation, as well as biological examples. We implemented a publicly available web server for visualizing and investigating the isoform-level network (http://guanlab.ccmb.med.umich.edu/isoformnetwork/).

The key element of multiple instance learning is to identify the functionally related isoform pairs (‘witnesses’) of the positive gene pairs. Due to the extremely large number of possible assignment of witnesses, the optimum solution is often sought through an iterative approach [46]. In previously introduced MIL algorithms such as mi-SVM and MI-SVM [46], the “witness” initialization was done by (1) randomly choosing one isoform pair or (2) treating all isoform pairs of a functionally related gene pair bag as “witnesses”, thus inevitably introducing a lot of false positives. To rationalize the initialization of “witness”, we proposed a novel single instance bag-based initialization method (SIB-MIL), and compared its performance against other variants of MIL in the context of the functional network prediction. What distinguishes SIB-MIL from other MIL methods is that, in the first iteration, a model is built using only those bags that contain only a single instance (**Materials and methods**). This was possible since 85% of the genes only have one confirmed splice variant in the NCBI mouse gene annotation database. We found our new SIB-MIL method works well in this context and achieves 5-30% improvement over previous methods (**Figure S2**).

### Simulation study shows that SIB-MIL algorithm accurately predicts functional relationships at the isoform level

Since no isoform-level gold standard is available for the real data, we first tested the performance of the SIB-MIL algorithm using simulated data under different scenarios. This simulation was carried out assuming that we only have 50 genomic datasets, which would result in a very conservative estimation of our real performance. Two parameters were tested in our simulation study: (1) the <u>discriminativeness</u> of the input data, represented by the mean difference (MD, see Figure 2A–C),) of the values between the population of functionally related isoform pairs and the population of functionally unrelated isoform pairs and (2) the multi-isoform gene ratio (MGR), defined as the ratio of multi-isoform genes to the total number of genes (see **Materials and methods**). For each simulation, we randomly partitioned gene pairs into two disjoint graphs as training and test set, respectively. Using disjoint graphs ensures non-contamination between the training and the test set, which is more appropriate than only separating gene pairs and allowing shared genes between the training and the test sets. We repeated the partitioning 20 times and thus tested our method on 20 randomly generated test sets. For each partition, we built a Bayesian classifier model using SIB-MIL and predicted a probabilistic functional relationship score for each isoform pair.

**Figure 2.**
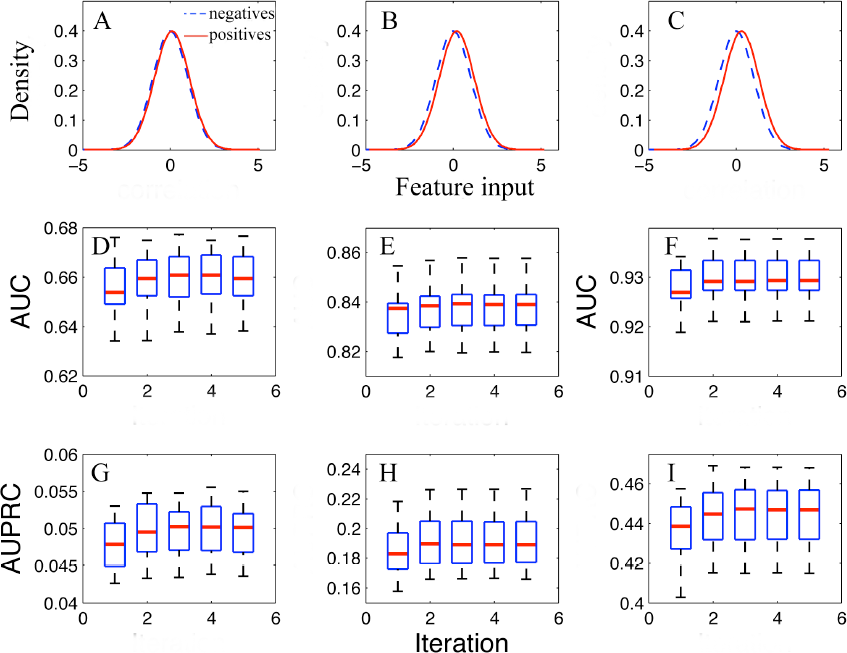
Predictive performance of the SIB-MIL algorithm on the simulated data at different values of mean difference of feature inputs between functional related (positives) and unrelated (negatives) isoform pairs. The multi-isoform ratio was fixed at 0.3. For both the positives and the negatives, the distribution of feature input was simulated with a normal distribution with standard deviation of 1. A-C shows the distribution of the feature inputs for the positives and the negatives, with mean difference = 0.1, 0.2 and 0.3 respectively. AUC and AUPRC figures below (D-I) show the corresponding classification performance at the isoform level. For all scenarios, the prediction performance of the SIB-MIL algorithm converges within 5 iterations. Prediction accuracy of SIB-MIL improves with increasing discriminativeness of the input feature data.

We first investigated the influence of the discriminativeness of the input data on the predictive performance of our SIB-MIL algorithm. We simulated that, for both the positive examples and the negative examples, the input feature values follow a normal distribution, with standard deviation equal to 1. Then, the mean difference (MD) values between the positives and the negatives can vary with the discriminativeness of the feature. With MGR fixed at 0.3, the prediction accuracy at the isoform pair level of SIB-MIL in terms of AUC with MD = 0.1, 0.2 and 0.3 are shown in Figure 2D, E and F. As expected, with increasing MD values the classification performance of SIB-MIL improved significantly. The median AUC on the 20 test sets for MD = 0.1, 0.2, 0.3 are 0.6595, 0.8389 and 0.9293, respectively. In addition, we also calculated the AUPRC (area under precision recall curve) and found that it also improves with increasing MD values (Figure 2G–I). This observation suggests that our SIB-MIL algorithm works well with input genomic data of very weak (MD = 0.1 or 0.2) discriminative power.

We further looked at how the predictive performance of our SIB-MIL algorithm will change with the multi-isoform gene ratio. To this end, we fixed the value of the mean difference of input data to be 0.2. The prediction accuracy at the isoform pair level of SIB-MIL in terms of AUC at MGR = 0.2, 0.3 and 0.5 are 0.8315, 0.8389 and 0.8275, respectively (Figure 3A, B and C). In addition, we also calculated the AUPRC (Figure 3D–F). Interestingly, we found that the performance gain of the converged model over the model at the first iteration increases with the fraction of multi-isoform genes. These gains for MGR = 0.2, 0.3, 0.5 are 0.0019, 0.0030 and 0.0124 for AUC, respectively, showing that SIB-MIL leads to better performance gain for more difficult situations. Overall, this shows that our algorithm is robust against the percentage of multi-isoform genes among all genes, and will remain to be applicable when new alternatively spliced variants are identified and verified.

**Figure 3.**
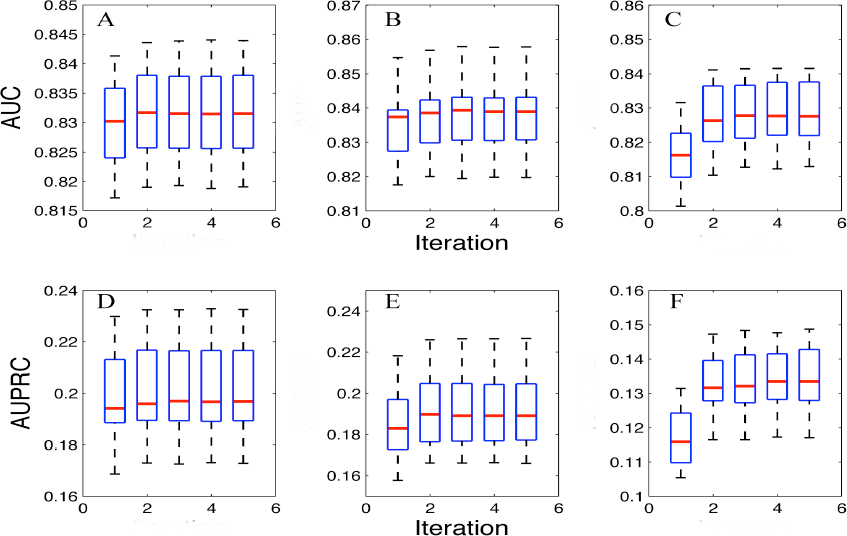
Predictive performance of the SIB-MIL algorithm on the simulated data at different multi-isoform gene ratios (MGR). The mean difference of the input features between functional related and unrelated isoform pairs was fixed at 0.2 (see Figure 2B). The MGR in A/D, B/E and C/F are 0.2, 0.3 and 0.5, respectively. For each setting (each plot), it can be seen that the prediction performance of the SIB-MIL algorithm converges within 5 iterations.

To investigate the combinatory effects of MD and MGR, we analyzed all the 9 simulations as mentioned above using SIB-MIL and found that for assigning isoform pair-level labels, SIB-MIL is robust to variations of the input data accuracy as well as the fraction of multi-isoform genes (**Figure S3-S5**).

### Modeling and validating the isoform-level functional relationship network for the mouse

To build a isoform-level functional network for the mouse, we calculated isoform-level pair-wise genomic features using multiple types of data sources, including RNA-seq, exon array, protein docking scores and pseudo-amino acid compositions (**Materials and Methods**). These pair-wise features may reflect and support the co-functionality between isoforms. We then generated gene-level gold standard from the genes co-annotated to a specific Gene Ontology term [29], or a BioCyc [31] or KEGG pathway [30] **(Materials and Methods)**. Using SIB-MIL, we built a genome-wide isoform-level network for the mouse by integrating the gene-level gold standard functionally related gene pairs and the isoform-level genomic features.

To computationally evaluate the effectiveness of our approach, we assigned each gene pair a score as the maximum probability of all its isoform pairs, under the assumption that co-functionality of a gene pair must be carried out by at least one of its isoform pairs. To test the predictive performance of SIB-MIL, we randomly generated 20 pairs of disjoint training sets and test sets for cross-validation (**Materials and Methods**). We found that our SIB-MIL method converges fast and achieves significantly higher AUC and AUPRC compared to the networks generated at the first iteration (Figure 4).

**Figure 4.**
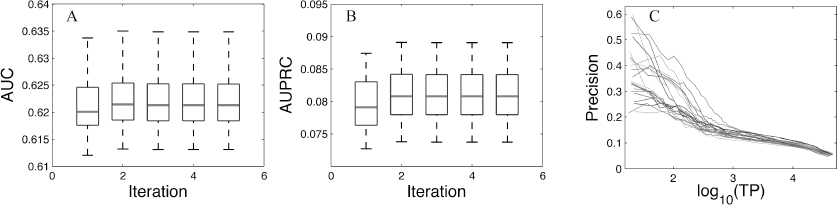
Performance of SIB-MIL for the mouse functional relationship network. This figure shows the results from 20 cross-validations. For each gene pair, the maximum value of all its isoform pairs was taken as its predicted value. The performance in terms of AUC (A) and AUPRC (B) was improved and converged within 5 iterations..

We next evaluated our predictions against several public databases, including protein-protein interaction [52–58], MSigDB gene sets [59] and Reactome pathways [41] (Table 1). Although these databases are not orthogonal to each other, they together provide a largely comprehensive source of both physical and functional interaction data. Because we only use the novel predictions (not recorded in GO, KEGG or BioCyc) for evaluation, these databases provide a rich resource to test the performance of novel predictions at least at the gene level. Again, we assigned the maximal probability of all its isoform pairs to each gene pair. We identified a list of 680,624 gene pairs with probability higher than 0.95 (**Table S1**). This represented a set of functionally highly related gene pairs that were newly predicted by our SIG-MIL algorithm. We found 36.0% of our top predicted gene pairs (244,957 gene pairs) are supported by co-annotation to the same biological process/pathway or having physical/genetic interactions, which is significant (*p* < 0.000001) compared to 7.0% of a set of randomly generated gene pairs. This result further supports the high precision of our network prediction model.

**Table 1.**
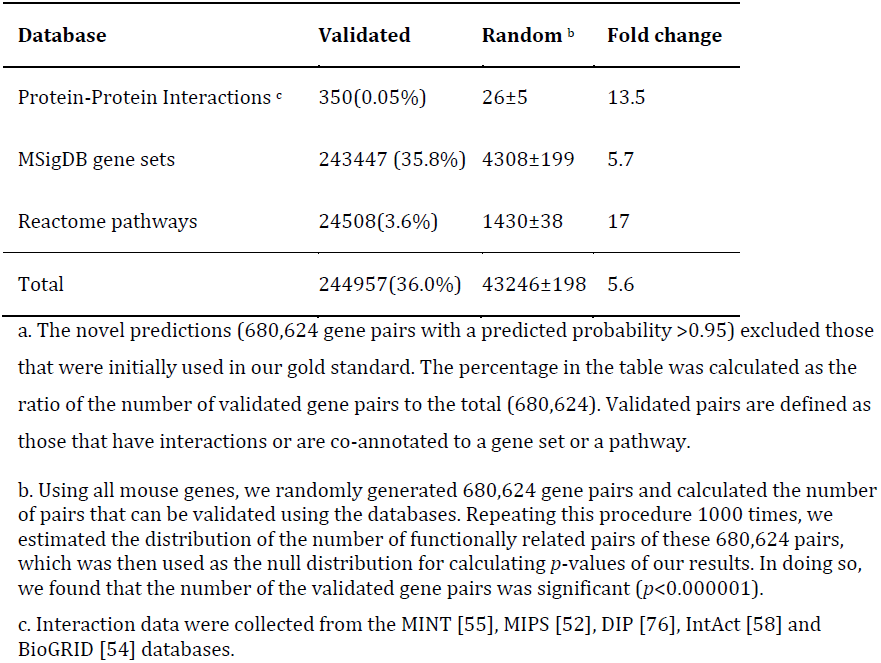
Validation results of our novel predictions of highly related gene pairs against pathway and interaction databases a.

### The isoform-level network provides a high-resolution map of functional relationships

As discussed in the Introduction, one major limitation of traditional gene-level functional relationship networks is that they consider a gene as a single entity and hence are not able to model functional connections for alternatively spliced isoforms. They treat a mixture of isoforms as a homogeneous entity. We found that our isoform-level functional relationship network for the mouse is capable of identifying differential functional relationships between isoform pairs belonging to the same gene pair. For example, the local gene-level network of *Ptbp1* reveals the functional relationship between any two genes as a single probability (Figure 5A**, left panel**). In fact, many genes in this network have multiple isoforms. For example, both *Ptbp1* and *Banf1* have two alternatively spliced isoforms. The functional relationship between *Ptbp1* and *Banf1* in our isoform-level network can therefore be dissected into four functional linkages corresponding to four possible isoform pairs (Figure 5A**, right panel**). Among the four isoform pairs, the functional relationship of the isoform pair [*NM_008956.2*, *NM_011793.2*] is predicted to be 0.999, whereas the probabilities of the other three isoform pairs are much lower---0.233, 0.084, and 0.045, respectively. These predictions are indeed reflected by the biology of these isoforms. The polypyrimidine tract binding protein (PTB), also known as *Ptbp1* or hnRNPI, is a key factor in RNA metabolism, and mRNA splicing events [60,61]. The translated proteins of its isoforms (NM_008956.2 and NM_001077363.1) have four quasi-RNA recognition motifs (RRM1-4) that bind to RNA to enable splicing mechanisms. The only difference between the proteins is that the translated form of NM_001077363.1 is 26 amino acids longer, which causes a shift in the positions of RRM3 and RRM4 motifs compared to that of the translated protein form of NM_008956.2 [62]. According to the isoform network predictions, *Banf1* was predicted as a key interactor of *Ptbp1*. *Banf1* isoform NM_011793.2 was predicted to interact with the shorter *Ptbp1* isoform NM_008956.2 with a probability of 0.999. The Gene Ontology enrichment analyses show that both the isoforms of *Banf1* as involved in cell cycle and cell division processes with a p-value smaller than 0.01. This is consistent with previously published reports that support the important roles of *Banf1* in chromatin structure formation, cell division and gene regulation [63–65].

**Figure 5.**
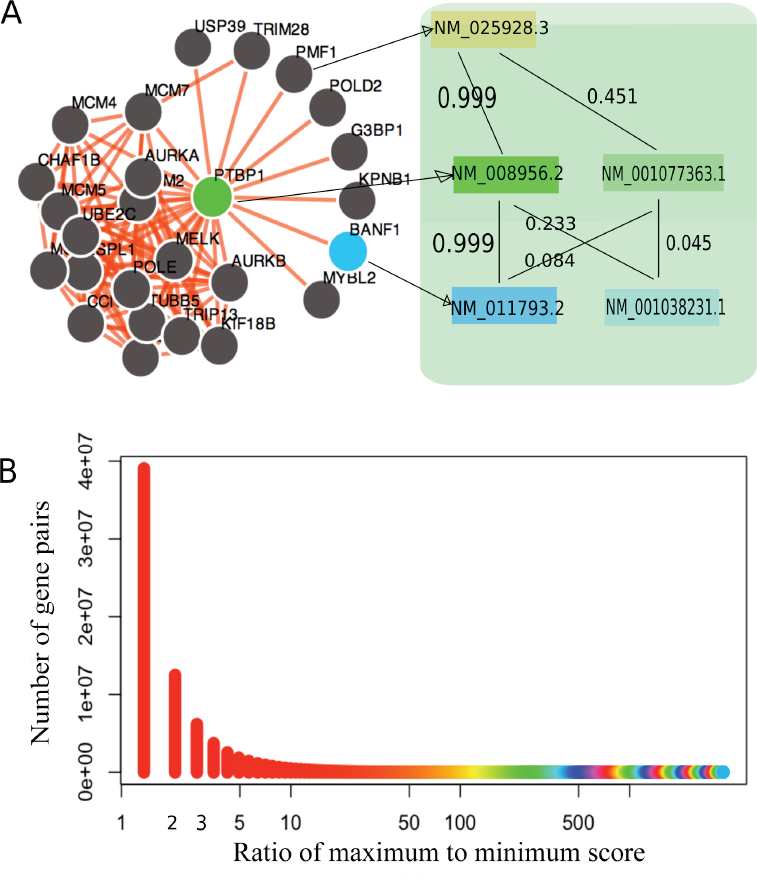
The isoform-level network reveals a high-resolution map of gene-gene connections. A. The left panel displays a traditional gene network of *Ptpb1* where there is only one functional linkage between each gene pair. Such a network is of low resolution since it considers a gene as a single entity and therefore is not able to describe functional relationships between isoform pairs of multi-isoform genes such as *Ptbp1*. Taking a gene pair [*Ptbp1* (with 2 isoforms), *Banf1* (with 2 isoforms)] as an example, its gene-level functional relationship is dissected into 4 isoform-level linkages among which the isoform pair [NM_008956.2, NM_011793.2] is most likely to be functionally related since its predicted probability is 0.999, which is significantly higher than that of the other three isoform pairs. As another example, the predicted isoform-level functional relationship scores between *Ptbp1* and *Pmf1* are also different. B. For each gene pair, we calculated the ratio of the maximum to the minimum probability among all isoform pairs. Shown here is the distribution of this ratio of 75,512,782 gene pairs. 25% of the gene pairs have an this value larger than 3.

On the other hand, it seems like the RNA splicing functionality of *Ptbp1* could be mainly through the longer protein product of NM_001077363.1 as the Gene Ontology analyses showed RNA processing, RNA metabolism and gene expression as the enriched functions for this isoform (Table 2). The shift in the relative RRM3 and RRM4 positions due to the absence of 26 amino acids in the translated protein of shorter *Ptbp1* isoform NM_008956.2 may have an influence on its role in RNA splicing [62]. This prediction is indeed supported by Wollerton et al [66], which experimentally showed that the longer variant of *Ptbp1* which has the additional 26 amino acids has more repressive effect on the splicing events compared to the other PTBP1 isoforms. Vitali *et al* reported the importance of the relative positions and tight interactions between RRM3 and RRM4 motifs in RNA-splicing [67], which is consistent with the different probabilities that we predicted for the interactions between the isoforms of these two genes.

**Table 2.** A subset of GO terms significantly enriched in the local isoform network of NM_008956.2 and NM_001077363.1 of *Ptbp1*.

| Isoform | GO term ID | GO term Name | *p*-value |
| --- | --- | --- | --- |
| NM_008956.2 | GO:0000278 | Mitotic cell cycle | 4.81E-17 |
|  | GO:0051301 | Cell division | 7.52E-14 |
|  | GO:0006270 | DNA replication initiation | 1.25E-10 |
|  | GO:0007599 | hemostasis | 2.69E-02 |
|  | GO:0033205 | cell cycle cytokinesis | 1.28E-02 |
|  | GO:0034501 | protein localization to kinetochore | 4.50E-02 |
| NM_001077363.1 | GO:0006396 | RNA processing | 7.30E-03 |
|  | GO:0016070 | RNA metabolic process | 1.33E-03 |
|  | GO:0010467 | gene expression | 2.82E-02 |

Such disparities of connections between isoform pairs of the same gene pair are prevalent. To quantify such difference, we calculated the ratio of the maximum to the minimum of the predicted probability among all isoform pairs of a given gene pair, respectively. For example, this ratio for the *Ptbp1* and *Banf1* gene pair is calculated as 0.999/0.045 = 22.20 where 0.999 and 0.045 are the maximum and minimum score between this gene pair, respectively (Figure 5A). From the whole isoform-level network of mouse, we found that this ratio spans a wide range from 1.0 (no difference) to more than 3500 (3500 fold difference) (Figure 5B). Notably, 25% of these gene pairs have a fold change value larger than 3.0, indicating that a high proportion of the gene pairs are functionally highly differentiated at the isoform level. These results suggest that difference is prevalent across isoform pairs coming from the same gene pair and that our isoform network can reveal such variations.

### The isoform-level network reveals functional diversity of different isoforms of the same gene

It is known that proteins encoded by isoforms of the same gene can carry out different and even opposite biological functions, such as pro-apoptotic versus anti-apoptotic actions of bclx-L vs bclx-S and of caspase 3 (L vs S) and transcriptional activation versus transcriptional repression for odd-skipped 2 [62]. Investigating and revealing the functional diversity of the same gene achieved by alternative splicing is pivotal to biology. Because of its high resolution, our isoform network has the ability to reveal such functional diversity.

To systematically examine the functional diversity represented at the network level, for each of the 3447 validated multi-isoform genes in the NCBI database, we compared the local networks (with the top 25 neighbors) between all its possible isoform pairs and counted the number of shared functionally related neighbors (**Figure S6**). We found that the minimum, mean and maximum numbers of shared neighbors are 0, 4 and 24, respectively. This statistics of isoforms’ neighborhood indicates that many isoforms of the same gene have different functional connections and may participate in different biological processes. For example, *Anxa6* has two alternatively spliced isoforms: NM_001110211.1 and NM_013472.4. Both isoforms have the same N- and C-termini, but the former encodes a shorter protein by six amino acids (525-530) due to the lack of an alternate in-frame exon compared to the latter. Using our web server, we identified the local networks of these two isoforms (Figure 6). Their local networks share 13 out of 25 neighboring isoforms, indicating a diversified functional relationship map of these two isoforms despite similar structures. To further reveal the functional differences of the genes in the two local networks, we performed Gene Ontology (biological process terms only) enrichment analysis using the top connected genes in each network (Table 3). We found that, while sharing the same GO terms (such as GO:0006944 cellular membrane fusion and GO:0061025 membrane fusion), the two isoform networks are also enriched for genes annotated to different GO biological processes. The isoform network of NM_001110211.1 is enriched for genes associated to vesicle fusion (GO:0006906, *p* = 0.0096), organelle fusion (GO:0048284, *p* = 0.044), and amino acid activation (GO:0043038, *p* = 0.0262), whereas the isoform network of NM_013472.4 is enriched for genes related to regulation of cell shape (GO:0008360, *p* = 0.0135). These disparate enriched functions strongly support the functional diversity of the two isoforms of the *Anxa6* gene. Our computational modeling of the folding and conformation of the two isoforms shows a striking difference in likelihood of phosphorylation in the Thr-Pro-Ser (535-537 vs 529-531) sequence [62]. In addition, the alternative splice isoforms of the *Anxa6* gene have been reported to have functional differences on catecholamine secretion [68], which is consistent with our functional enrichment analysis related to the vesicle fusion and organelle fusion (Table 3). These results suggest that our isoform-level network is able to reveal functional diversity of different isoforms of the same gene and could therefore become a promising tool for investigating gene functions at the refined isoform level. To facilitate new isoform function discovery based on our network, we included this enrichment analysis for all local networks of individual isoforms in our interactive website.

**Figure 6.**
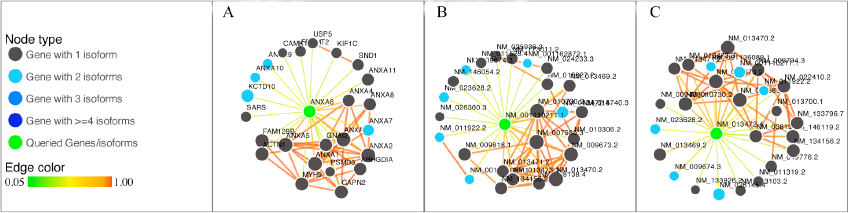
The gene-level functional relationship network of the *Anxa6* gene (A) and the isoform-level networks of its two isoforms: NM_001110211.1 (B) and NM_013472.4 (C). The top 25 neighbors of the query gene are shown and the probability threshold for between-neighbor links is set to 0.99. The local networks of the two isoforms reveal different connections reflecting their respective functional roles.

**Table 3.** GO terms significantly enriched in the local isoform network of NM_001110211.1 and NM_013472.4 of *Anxa6 \**.

| Isoform | GO term ID | GO term Name | p-value |
| --- | --- | --- | --- |
| NM_001110211.1 | GO:0006944 | cellular membrane fusion | 2.75E-03 |
|  | GO:0061025 | membrane fusion | 3.59E-03 |
|  | GO:0031340 | positive regulation of vesicle fusion | 1.65E-02 |
|  | GO:0007599 | hemostasis | 2.69E-02 |
|  | **GO:0006906** | **vesicle fusion** | **9.66E-03** |
|  | **GO:0006418** | **tRNA aminoacylation for protein translation** | **2.11E-02** |
|  | **GO:0043039** | **tRNA aminoacylation** | **2.62E-02** |
|  | **GO:0043038** | **amino acid activation** | **2.62E-02** |
|  | **GO:0048284** | **organelle fusion** | **4.39E-02** |
|  | **GO:0031338** | **regulation of vesicle fusion** | **4.61E-02** |
| NM_013472.4 | GO:0006944 | cellular membrane fusion | 4.11E-03 |
|  | GO:0061025 | membrane fusion | 5.38E-03 |
|  | GO:0031340 | positive regulation of vesicle fusion | 2.09E-02 |
|  | GO:0007599 | hemostasis | 4.01E-02 |
|  | **GO:0008360** | **regulation of cell shape** | **1.35E-02** |
|  | **GO:0044699** | **single-organism process** | **4.20E-02** |

\*, the uniquely enriched GO terms are in **Bold**.

### Webserver

We implemented a webserver to search and visualize the mouse functional network at the isoform level using Mysql, PHP as well as javascript. It is publicly available at (http://guanlab.ccmb.med.umich.edu/isoformnetwork).

## Discussion

In the past decade, significant efforts have been devoted to model functional relationship networks at the gene level, including global, tissue-specific and biological process-specific networks [1–25]. These networks have been established in several model organisms to predict biological functions, pathway components, and phenotype-associated genes [1,18,19,21,32,33]. However, all these works considered a gene as a single entity without differentiating functional relationships between multiple isoform pairs within a gene pair. Building functional networks at the isoform level promises a high-resolution map of the traditional functional gene networks. However, due to the lack of isoform-level gold-standard functionally related pairs, establishing networks at the isoform level is a challenging problem.

Inspired by our recent approach in assigning isoform-level functions [49], we developed a novel multiple instance learning algorithm and integrated it with the Bayesian network, which has been widely used in the network modeling field, to build an isoform-level functional relationship network for the mouse. Our simulation studies showed that our method can accurately predict the functional relationships at the isoform level. We further built a genome-wide isoform network for the mouse, and the evaluation results based on gene-level prediction accuracy, the validation against existing databases and the validations using individual isoforms again demonstrated that our method is successful in predicting functional relationships at both the isoform- and the gene-level.

Despite these advantages, our isoform-level network has the following limitations. First, our gold standard includes only protein-coding genes. Therefore, the predictions of non-coding pairs made by our model, though informative, may be biased. Second, what we have built is a global isoform-level network without differentiating functional relationships in different tissues. Tissue-specific functional relationship network at the gene level is an emerging field right now [1]. Building tissue-specific networks at the isoform level will be of great value to reveal the spatial dynamics of functional interactions between isoforms, which is beyond the scope of this study.

Overall, our isoform-level network represents a novel approach to probe functional relationships at the isoform level, thus providing a higher resolution view compared to traditional gene-level networks. Investigating isoform-level networks would help deepen our understanding of gene functions and functional relationships, and may provide useful information on diseases caused by alternative splicing. We expect that isoform-level networks will find wide applications in genomic and biomedical applications, and that the current gene-centered network modeling approach will be expanded to a more refined isoform level.

## Materials and Methods

### Single-instance bag based multiple instance learning (SIB-MIL) algorithm

The key point of multiple instance learning (MIL) [43,46,50,51,69] is to iteratively identify the functionally related isoform pairs (‘witnesses’) of positive gene pairs. In previously developed MIL algorithms such as mi-SVM and MI-SVM [46], the witness initialization was done by either randomly choosing one isoform pair or treating all isoform pairs of a functionally related gene pair (a positive bag) as witnesses. Thus, these methods could inevitably introduce false positives. To make the initialization of witnesses more rational, we proposed a novel single-instance-bag-based multiple instance learning (SIB-MIL), which is described below.

Without loss of generality, the *i*th gene pair containing *m* isoform pairs is denoted by *X*_*i*_ ={**x***_i_*_1_, **x**_*i*2_…**x***_im_*}with **x***_ij,_ j = 1,2…m* denoting the *j*th isoform pair of the *i*th gene pair. We assign the class label of the *i*th gene pair, denoted as *y*_*i*_, based on our hypotheses that for a positive gene pair, **<u>at least one</u>** of its isoform pairs is functionally related; if a gene pair is negative, **<u>none</u>** of its isoform pairs should be functionally related, which can be mathematically expressed as follows:

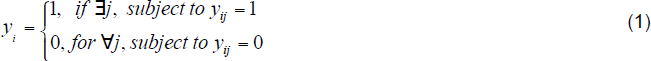

where *y*_*i*_ indicates the label of the *j*th isoform pairs of the *i*th gene pair. We refer to those positive instances (functionally related isoform pairs) of a positive bag (positive gene pair) as the witnesses. The iterative solution is detailed as the following:

(1) Initialization: because there is no existing isoform-level gold standard, a set of isoform pairs in positive gene pair bags need to be selected as witnesses to build an initial model. Treating all isoform pairs as witnesses or randomly choosing one pair as the witness is likely to introduce false positives. To overcome this issue, we chose only those bags in which both genes are single-isoform genes to construct the initial set of -witnesses- at the isoform level. In this particular setting, we have sufficient number of single-isoform genes to construct the initial set of -witnesses-. This approach was motivated by the fact that the isoform pair in these single-instance positive gene pair bags must be positive and thus will not introduce any false positives. Therefore, all the instances in these single-instance gene pair bags are labeled as Class 1 (functionally related). All instances in negative bags are labeled as Class 0 (functionally unrelated). In the following simulation studies, we compared this SIB-MIL method with several previously established methods, and showed the superior performance of SIB-MIL (**Figure S2**).
(2) The loop:

(2.1) Model building: using the current witness set and the negative isoform pairs, we build a new Bayesian network classifier that will be used to re-assign a probability score to all instances in the original training set.
(2.2) Witness updating: For each positive bag, reselect the instance with the maximum probability score as the “witness” and label it as Class 1. Also for each negative gene pair bags, only the highest scored instance was chosen and labeled as Class 0 for model building. The reason to choose the highest scored instances in negative bags is that a classifier is expected to perform well if it can correctly classify the most difficult examples.
(3) Stop criteria and final predictions: The iteration is stopped when cross-validation performance does not change any more. The final classifier, made at the instance (isoform pair) level, will be used to predict the final network. Each isoform pair will be assigned with a probability to be functionally related. At the gene-pair bag level, the score of each bag is assigned to be the maximum of all scores of its instances.

In each iteration, we used a Bayesian network classifier, as previously described in the work [1,2,8,14] as the base learner in our SIB-MIL implementation. Briefly, Each isoform pair can be represented by an *n*-dimensional feature vector (*E*_1_,*E*_2_…*E*_n_). With the Bayesian classifier, the probability that an isoform pair belonging to the positive class can be calculated using the following formula:

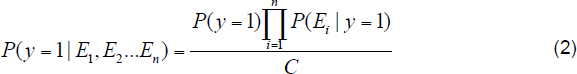

Where *P*(*y* = 1) is the prior probability for a sample to be positive, *P(E_i_|y = 1)*, i = 1,2,…,n, is the probability of the *i*th feature with the observed value, given that the isoform pair is functionally related and *C* is a constant normalization factor.

### Simulation studies

We simulated a series of scenarios to examine the ability of our algorithm to predict functional relationships at the isoform level. In this simulation study, we set the number of genes to 5000 and as in the real NCBI database, a gene may contain one or several isoforms. The number of positives (functionally related gene pairs) was set to 10, 000. The number of negatives (functionally unrelated gene pairs) was set to 19 times that of positives based on our previous study [1]. The number of isoform-pair level features was set to 50.

We focused on examining the effects of two factors on the performance of our algorithm. 1, the discriminativeness of the feature data and 2, the fraction of the multi-isoform genes among all genes. For each feature, we simulated that the distributions of the positive examples and the negative examples both follow normal distributions with a standard deviation of 1. Then, the discriminativeness of features is controlled by the Mean Difference (MD) between the population of functionally related isoform pairs and the population of functionally unrelated isoform pairs (Figure 2). In our study, three MD values, *i.e.* 0.1, 0.2 and 0.3, were tested. For the second factor, based on the RefSeq, which is a validated database of genes and isoform annotation, a gene may contain a single or multiple isoforms. So, the ratio of multi-isoform genes to the total number of genes (MGR) should be considered. For this ratio, we tested three values: 0.2, 0.3 and 0.5. For example, MGR = 0.5 means that half of the genes are multi-isoform genes. The source codes in Perl for the simulating data are available at: http://guanlab.ccmb.med.umich.edu/isoformnetwork/download.php.

### Collecting and pre-processing isoform-level genomic data

We had in total 164 isoform-level features: 41 from RNA-seq data, 121 from exon array, 1 from pseudo-amino acid composition and 1 from protein-docking score data. Details for processing these four types of data are described below. Protein domain data was excluded due to direct Gene Ontology annotation transfer from domain information in mouse.

#### RNA-seq

We downloaded 117 mouse RNA-seq datasets (corresponding to 811 experiments) from the NCBI sequence read archive (SRA) [70] on May 1, 2012, which cover a wide range of experimental conditions and different tissues. For each RNA-seq experiment, we used the TopHat (v2.0.051) [34,71] to align the reads against the *Mus Musculus* reference genome from the NCBI gene build (version 37.2). Then, the resulting mapped read files together with the corresponding transcript annotation files were processed by Cufflinks (v2.0.0) [34] to calculate the relative abundance of the transcripts in terms of FPKM (Fragments Per Kilobase of exon per Million fragments). We removed those experiments with less than 10 million reads or covering less than 50% of the genes. In addition, to calculate correlations, those datasets with fewer than 4 experiments were also removed. In doing so, we finally obtained 41 datasets including 386 experiments (**Table S2** for a summary of these experiments). Within each dataset, we further removed those transcripts with more than 50% missing values to ensure the accuracy of expression correlation estimation. FPKM values were log-2 transformed as they were treated in Cuffdiff 2. We calculated Pearson correlation coefficients, denoted by *ρ*, between all possible transcript pairs for each dataset, followed by normalization using Fisher’s z-transformation [72] to allow comparison between different datasets:

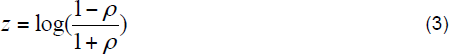

This *z*-transformed correlation will be used as isoform-level features when building the Bayesian classifiers through the multiple instance learning approach.

#### Exon array data

121 mouse exon array (Affymetrix Exon 1.0 ST array) datasets of the mouse were downloaded from the NCBI GEO (Gene Expression Omnibus) database (**Table S3** for a summary of these experiments). Each dataset includes at least 4 experiments. We calculated the expression of transcripts by utilizing the R package MEAP (version 2.0.1) [73]. Then, The Pearson correlation coefficient between each pair of transcripts was computed and normalized using the Formula (3). These correlations will be used as the feature inputs of our SIB-MIL algorithm.

#### Pseudo-amino acid composition

Pseudo-amino acid composition (pseAAC) is a descriptor that characterizes the standard amino acid composition (AAC) as well as the pseudo-AAC by taking the sequence information of a protein into account [74]. Here, we calculated pseudo-amino acid composition (pseAAC) data for the protein-coding isoforms. The number of pseudo components was set to be 20. So each protein sequence was characterized by a 40 dimensional vector (composition of 20 natural amino acids plus 20 pseudo AACs). Then we calculated Fisher’s *z*-transformed Pearson correlation between isoform pairs as the feature data.

#### Protein docking score

We computed and derived a quantitative physical interaction score for each isoform pair using the SPRING algorithm [75]. Briefly, SPRING is a template-based algorithm for protein-protein structure prediction. It first threads one chain of the protein complex through the PDB (Protein Data Bank) library with the binding partners retrieved from the original oligomer entries. The complex models associated with another chain are deduced from a pre-calculated look-up table, with the best orientation selected by the SPRING-score, which is a combination of threading Z-score, interface contacts, and TM-align match between monomer-to-dimer templates.

These four types of feature datasets together provide a largely comprehensive characterization of isoform pairs. They cover information from sequence, expression, physical interaction as well as amino acid composition. To remove potential uninformative feature datasets, we evaluated each of these 169 datasets against a gold standard, in which all isoform pairs of a positive gene pair are initialized as positives, and removed those datasets with AUC lower than 0.51. Finally, 65 feature datasets were retained for building the final isoform-level network (**Table S4**).

### Gene-level gold standard functionally related pairs

We constructed a gene-level gold standard of functionally related pairs using the Gene Ontology (GO) [29], KEGG [30], and BioCyc [31] databases. Gene Ontology is organized into a hierarchy where broader terms have more genes annotated to each but represent non-specific biological functions, while specific terms have few genes annotated to each. Some of the GO terms are too broad to be experimentally tested, such as ‘metabolic process’, and gene pairs co-annotated to such terms cannot be considered as truly functionally related. We therefore used a list of Gene Ontology terms voted by the biologists, which represent a wide spectrum of experimentally testable biological processes [42], and excluded the terms with more than 300 annotated genes. A pair of genes is considered to be functionally related if they are co-annotated to the same specific biological process or involved in the same biological pathway as defined by KEGG or BioCyc. Such a gene pair is defined as positive. In total, we obtained 675,124 positive gene pairs.

Unlike positive pairs, there is no database that defines two genes as functionally not related. Consistent with previous works in this field [1,3], we used random pairs as negatives and fixed a ratio of negatives to positives as 19:1. This ratio serves as the initial prior, and is fixed throughout the iterative process to ensure a consistent prior for Bayesian network prediction in each step.

## Acknowledgement

We thank Thomas Werner for insightful discussion. This work is supported by NIH 1R21NS082212-01, EU-FP VII Systems Biology of Rare Disease and NIH University of Michigan O’Brien Kidney Translational Core Center.

## Author Contributions

Y.G. and H.D.L. conceived the research. H.D.L. and Y.G. developed the core algorithm and performed computational analysis, H.D.L., Y.G. and G.S.O wrote the paper. R.E. processed the raw RNA-seq data. R. M. provided support for *Ptbp1* and *Anxa6*. A.G. and Y.Z. computed the protein-protein docking score.

## Supplementary materials

**Table S1**. Gene pairs with probability > 0.95 predicted by the mouse isoform-level network.

**Table S2**. The 41 mouse RNA-seq datasets downloaded from the Short Read Archive (SRA) database.

**Table S3**. The 121 mouse Exon array datasets downloaded from Gene Expression Omnibus (GEO) database.

**Table S4**. The selected datasets for building the mouse isoform-level network.

**Figure S1. Formulating the isoform-level network prediction into a multiple instance learning (MIL) problem. A**. Illustration of a functionally related gene pair (a positive bag), gene I with 3 isoforms and gene II with 2 isoforms. In this case, there are in total 6 possible isoform pairs. Among these, two isoform pairs are functionally related (solid red line), whereas the other 4 isoform pairs have no functional relationship (dashed light blue line). **B.** Illustration of a functionally unrelated gene pair (a negative bag), gene III with 2 isoforms and gene IV wiith 2 isoforms. None of the isoform pairs between III and IV can be functionally related. **C.** In the traditional gene-level network prediction, a classification model can be established to distinguish the positive examples, defined as known functionally related gene pairs, against the negative examples (unrelated pairs). **D.** In the isoform-level network prediction, *i.e.*, MIL, gene pairs are considered as ‘bags’, each of which may contain one to many isoform pairs, defined as ‘instances’. A positive bag (a co-functional gene pair, green oval) must have at least one of its instances being functionally related, which are called ‘witnesses’ (pairs in red). All instances (isoform pairs) in a negative bag (an unrelated gene pair) must not be functionally related. A classifier is trained under the above constraints.

**Figure S2. Performance comparison of different MIL algorithms in terms of ROC curves computed on 20 randomly generated test sets**. Version A: the MI-SVM algorithm proposed in the work [46] where a randomly selected isoform pair from gene pair bag is used as “witness” in its first iteration. Version B: a test version of MIL developed in our study whose initialization step is the same as that in Version A. From the second iteration, a subset of isoform pairs from negative gene pair bags were selected so as to keep the ratio of negative to positive isoform pairs the same as that in the first iteration. Version C: our proposed single-instance bag MIL (SIB-MIL).

**Figure S3.** Performance (in AUCs) of SIB-MIL on the simulated data, with 9 settings of MD and MGR values.

**Figure S4.** Performance (in AUPRCs) of SIB-MIL on the simulated data, with 9 settings of MD and MGR values.

**Figure S5.** Performance (in precision-recall curves) of SIB-MIL on the simulated data, with 9 settings of MD and MGR values.

**Figure S6.** The distribution of the number of shared interactors (out of the top 25 interactors) between any two isoforms of the multi-isoform genes.

